# Circadian proteomics reveal rampant tuning of post-transcriptional apparatus by *Chlamydomonas* clock

**DOI:** 10.1101/2023.07.28.550970

**Authors:** Dinesh Balasaheb Jadhav, Sougata Roy

## Abstract

Timing of biological processes enable organisms to sustain the diurnal fluctuations resulting from earth’s rotation. Circadian clocks execute this temporal regulation by modulating temporal expression of genes. Clock regulation of mRNAs was envisioned as the primary driver of daily rhythms. However, mRNA oscillations often don’t concur with the downstream protein oscillations. To assess the contribution from post-transcriptional processes, we quantitatively probed the *Chlamydomonas* proteome for two circadian cycles. Our study suggests rampant role of posttranscriptional processes in clock regulation of *Chlamydomonas* metabolism. We quantified >1000 proteins, half of which demonstrate circadian rhythms. Among these rhythmic proteins, >40% originate from non-rhythmic mRNAs and > 90% peak around midday or midnight. Accumulation rhythms of proteins rather than their encoding mRNAs shows extreme coordination. We uncovered new rhythms and accounted for physiological rhythms whose mechanistic details remained undocumented from earlier transcriptomic studies. We envisage our study will refine and enrich the evaluation of temporal metabolic processes in *Chlamydomonas.* Owing to *Chlamydomonas’s* unique phylogeny this study can lead to new insights into evolution of clock regulation across kingdoms.

## Introduction

Almost every organism inhabiting earth encounters temporal fluctuation in light and temperature over the 24-hour cycle. The organisms adapt by anticipating these temporal fluctuations and synchronizing their cellular activities to the ideal environmental time (Dunlap, 1999). This temporal regulation is accomplished by the cell-autonomous circadian clock, which can function even when the environmental cues are withdrawn (Aschoff, 1965). Consequences of this clock regulation are overt temporal rhythms in behaviour, physiology, and metabolism over the course of the 24 hour cycle (Takahashi, 2017b). This time-dependent cellular dynamics is achieved by the clocks’ ability to alter the expression of genes depending on the time of the day (Buhr & Takahashi, 2013). For a long time, the consensus relied on transcription to be the primary driver of this daily regulation (Papazyan *et al*, 2016). The discovery of rhythmically active promoters (Bell-Pedersen *et al*, 2005; Young & Kay, 2001), temporal regulation of promoter accessibility (Bell-Pedersen *et al*, 1996), and temporal dynamics of mRNA abundance (Doherty & Kay, 2010), all suggested that transcription would be the simplest way to relay the regulatory information to the downstream effector molecules. However, the discovery of the cyanobacterial circadian system shifted the paradigm (Cohen & Golden, 2015). Post-transcriptional mechanisms started gaining importance in temporal regulation of physiology and metabolism (Reddy & Rey, 2016). Indeed, this resulted in uncovering novel post-transcriptional strategies that are recruited by the clock to achieve this daily regulation (Edgar *et al*, 2012; Green, 2018; Kojima *et al*, 2011). The daily rhythms in physiology and metabolism can persist without transcription and this phenomenon is conserved across the eukaryotic kingdom (O-Neill & Reddy, 2011; Edgar *et al*, 2012; Johnson *et al*, 2017; Cohen & Golden, 2015; Tomita *et al*, 2005; Roy *et al*, 2014). Clock modulation of posttranscriptional processes significantly contributes to the temporal regulation of physiology and metabolism.

Although transcription has been the primary point of investigation regarding clock regulation, recent research suggests clock modulation can occur at multiple levels of gene expression. In *Lingulodinium polyedra,* the clock regulates protein synthesis and degradation rather than RNA synthesis (Morse *et al*, 1989) and transcriptome-wide analysis revealed no significant variation in mRNA abundance across diurnal and circadian cycles (Roy *et al*, 2014). In cyanobacteria phosphorylation and dephosphorylation of the clock protein is sufficient to generate the daily rhythms (Iwasaki *et al*, 2002). Studies in mammalian model systems also revealed that mRNA rhythms often do not translate to the downstream protein rhythms (Liu *et al*, 2016). For instance, studies with mammalian cells have shown that only 4.4% of rhythmic mRNA translates to rhythmic proteins (Mauvoisin *et al*, 2014). In murine liver (Robles *et al*, 2014) and suprachiasmatic nucleus (Chiang *et al*, 2014; Deery *et al*, 2009) 50% or more of the oscillating proteins exhibit corresponding oscillations in their parent mRNAs. The other half of proteins oscillate because of post-transcriptional regulation. Besides, 50% of the rhythmic mRNAs whose encoded proteins also oscillate showed significant mismatch in their respective phase distributions (Robles *et al*, 2014). Poor correlation between mRNA and protein rhythms were predominant in Drosophila S2 (Schneider 2) cells and the fungus *Neurospora crassa*, where almost 93% (Rey *et al*, 2018) and 41% (Hurley *et al*, 2018) of rhythmic proteins arise from non-rhythmic mRNAs respectively. These studies showed protein rhythms can arise independent of mRNA rhythms and peak accumulation time of proteins can differ substantially from its parent mRNAs. Thus, deducing the daily phase distribution of key cellular metabolic activities solely from the transcriptomic studies requires reconsideration. Rhythmic accumulation of proteins is equally important to understand the clock-controlled dynamics of cellular activities (Mauvoisin, 2019; Mauvoisin *et al*, 2017). In part, technological limitation of mass spectrometry (MS) is responsible for the lack of circadian proteomics studies. With the novel upgrades of the MS technology (Aebersold & Mann, 2016), it is now possible to assess the daily variation in physiology and metabolism by directly measuring the protein levels rather than inferring it from the mRNA levels.

Unicellular eukaryotic clock models are particularly interesting in understanding how clock regulate cellular metabolism, and physiology (Noordally & Millar, 2015), because they can easily synchronize to the light: dark cycles (Petersen *et al*, 2022; Matsuo & Ishiura, 2010; Ryo *et al*, 2015), they are clonal in nature and therefore amenable to biochemical studies (Salomé & Merchant, 2019), they provide a simple system with physiology analogous to multicellular organisms (Harris, 2001) and they lack the redundancy and complexity resulting due to intercellular interference. *Chlamydomonas reinhardti* is a unicellular chlorophyte (Harris, 2001; Mittag & Wagner, 2003; Sasso *et al*, 2018; Salomé & Merchant, 2019) with plant-like circadian system and many animal like features (Merchant *et al*, 2007). Owing to its unique relationship to the common ancestor of animals and plants (Brunner & Merrow, 2008) understanding *C. reinhardtii* circadian system could shed light to the evolution of circadian clocks and their mode of regulation across species. Plethora of known temporal rhythms such as starch metabolism (Ral *et al*, 2006) stickiness to glass (Straley & Bruce, 1979), nitrogen uptake (Byrne *et al*, 1992), cell division (Goto & Johnson, 1995), chemotaxis (Byrne *et al*, 1992), phototaxis (Johnson *et al*, 1991), UV-sensitivity (Nikaido & Johnson, 2000), etc are well-documented in *C. reinhardtii*. The *C. reinhardtii* phototaxis rhythm persists in space (Mittag *et al*, 2005; Mergenhagen & Mergenhagen, 1989), a study that finally validated that clocks are cell autonomous and can function without any geophysical cues. As yet, diurnal (Panchy *et al*, 2014; Strenkert *et al*, 2019) (Zones *et al*, 2015) and circadian (Kucho *et al*, 2005) transcriptomic studies in *C reinhardtii* formed the basis to infer the molecular correlates underlying the daily physiological rhythms and its phase distribution across the 24 hour cycle. Transcriptome-wide studies showed 80% to 85% of the measured mRNAs have daily rhythms (Strenkert *et al*, 2019). Although transcriptional rhythms are well studied in *C. reinhardtii*, a significant contribution of clock mediated regulation of cellular processes is also expected from the post-transcriptional mechanisms (Iliev *et al*, 2006). RNA-binding proteins (RBPs) play crucial roles in clock controlled biological processes (Mittag & Waltenberger, 1997; Mittag, 1996). In *C. reinhardtii*, CHLAMY1, a RBP, acts as a clock component and can repress circadian protein synthesis by directly binding to the 3’UTR of mRNAs (Mittag, 1996; Zhao *et al*, 2004; Waltenberger *et al*, 2001). Despite these findings there has been no comprehensive studies to account for the circadian dynamics of the proteome in *C. reinhardtii*, which are essential to complement the existing transcriptomic studies.

Due to technological impediments, earlier efforts to study the dynamics of circadian proteome in *C. reinhardtii* had limited scope (Wagner *et al*, 2004, 2005). Modern mass spectrometry technology allows reproducible measurement of proteomes across samples (Mauvoisin & Gachon, 2020). Sequential window acquisition of all theoretical mass spectra (SWATH-MS), a data-independent acquisition (DIA) approach (Sajic *et al*, 2015; Domon & Aebersold, 2010) that revolutionized the proteomics study by combining proteome coverage with consistent and reproducible quantification of the proteins (Collins *et al*, 2017, 2013). SWATH-MS can be a valuable technology for probing proteome wide circadian dynamics. We used SWATH-MS to study the circadian proteome of *C. reinhardtii*, to understand and estimate the clock involvement at the post-transcriptional level. We envisage that by quantifying and comparing protein intensity across the circadian cycle we can (1) uncover the clock controlled protein counterparts of the known physiological rhythms, (2) find novel physiological and metabolic pathways where clock regulation of proteins rather than mRNAs may have a role, (3) estimate the overall role of protein dynamics in circadian output and (4) find out the extent of concurrence between the rhythm of mRNAs and their encoded proteins. We were able to faithfully measure the intensity of 1080 proteins in two subsequent circadian cycles. Out of the total measured proteins, 48% appears to be rhythmic over two circadian cycles. Phase distribution analysis of cycling proteins yielded two major groups, about 56% were peaking during the day while around 44% proteins peaked at night. Functional categorization of the day and night peaking proteins allowed us to identify hundreds of enzymes associated to key cellular processes such as photosynthesis, light sensitivity, amino acid biosynthesis, redox metabolism, fatty-acid biosynthesis process, carbon fixation, glycolysis, oxidative phosphorylation, and tricarboxylic acid cycle. Peak phase of the proteins required to execute the above pathways were concurrently regulated. Our circadian proteome further uncovered novel rhythmic processes that were not uncovered in the earlier diurnal/circadian transcriptome studies. Our study also found widespread inconsistency in phase distribution of mRNAs and its encoded proteins. We found almost 42% of the cycling proteins are encoded from non-oscillating mRNAs. For the remaining 58%, where both mRNAs and proteins oscillate, 65% of the proteins have phase distribution that was significantly different from their parent mRNAs. Altogether, our results indicate a significant contribution of the rhythmic proteome in circadian regulation of physiology and metabolism and suggest extensive tuning of the post-transcriptional apparatus by *C. reinhardtii* clock.

## Results

### SWATH-MS captures the temporal rhythmic proteome of *C. reinhardtii*

*C. reinhardtii* cells were released in constant light (LL) after they were synchronized to the daily 12h:12h LD (light: dark) cycles for at least 5 days (Figure 1a). Samples in triplicate were collected from two independent circadian cycles at designated times (Figure 1a), proteins were purified and digested with trypsin to generate peptides, which were then sent to the mass spectrometry facility for data-independent acquisition in SWATH mode (Figure 1a). As a primary requirement of SWATH studies, we first generated the spectral library for *C. reinhardtii* using the data dependent MS approach (DDA). This library consists of 7681 spectra, 5335 distinct peptides, and 1506 proteins at a stringent 1% FDR. Using this library, we obtained quantitative information of 1269 proteins in total from the two cycles, which accounts for about 10% of the total annotated proteins in *C. reinhardtii*. However, we would like to point out that this is similar to the normal coverage obtained from any SWATH-MS studies conducted in other species (Collins *et al*, 2017; Basak *et al*, 2015; Chen *et al*, 2021). In this process, we generated the circadian spectral library of *C. reinhardtii* that can be a useful resource for future quantitative proteomics studies. Although the two-cycle replicates were collected and processed simultaneously, the mass spectrometry measurements of the 2 cycles were done days apart. Comparing the two cycles we found ~85% (1080) of the total 1269 proteins were common in both (Figure 1b). GO term analysis of our captured proteome has representations from different organelles and functional categories, which exhibit a faithful representation of the GO distribution from the total proteome (Supplementary Figure S1A). We also verified whether our captured proteome showed any bias based on its physio-chemical properties. It is evident from our analysis that when compared to the total proteome, the subset we captured is not biased to any physio-chemical properties (Supplementary Figure S1B). Consequently, we conclude that the proteome subset we captured is an unbiased representation of the total proteome. Therefore, we decided to proceed further with the 1080 common proteins obtained from the 2 cycles. We performed a published batch correction method COMBAT (Zhu *et al*, 2021) to remove any inconsistency due to non-biological factors within the two-cycle replicates (explained in detail in the M&M section).

**Figure 1:**
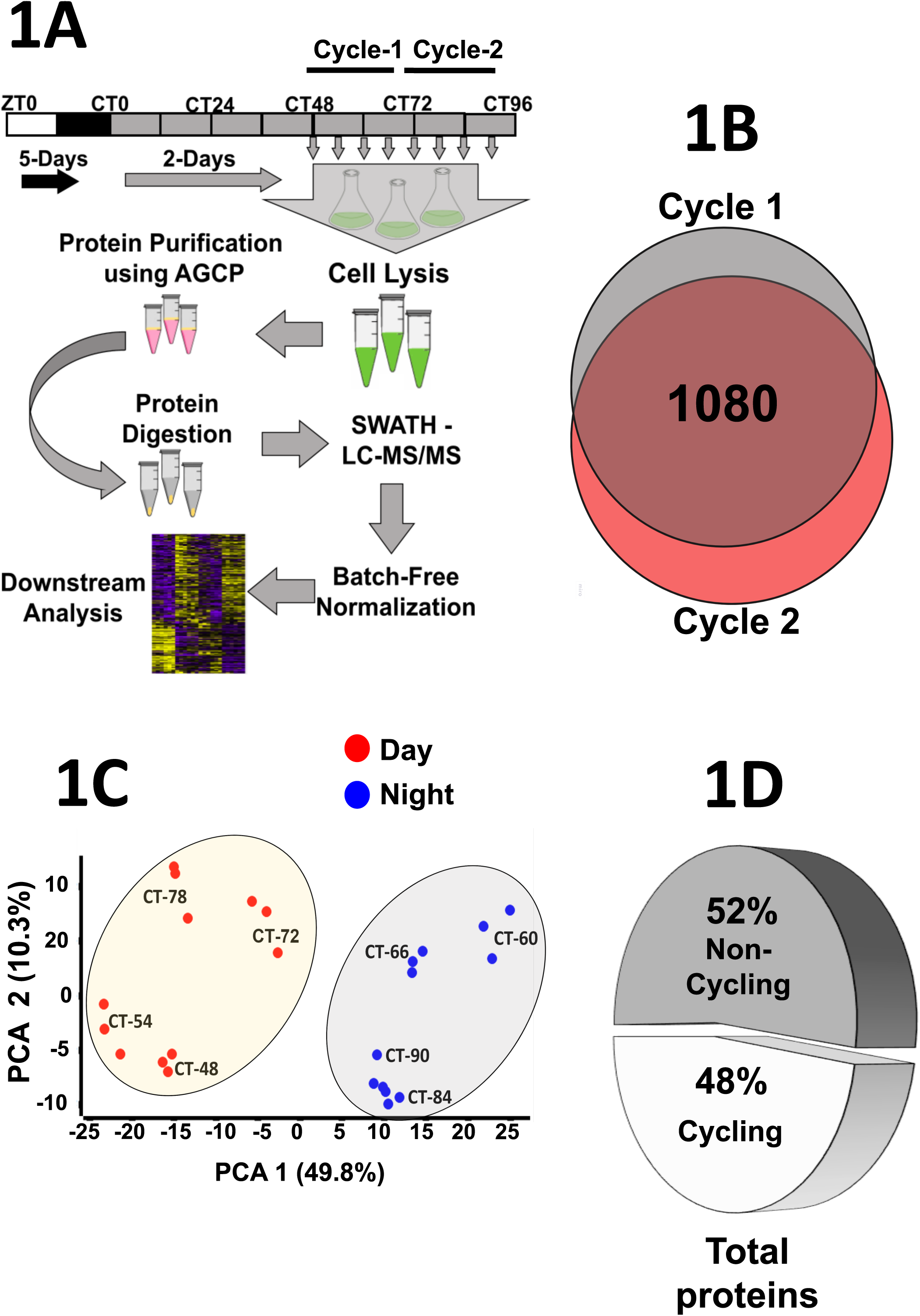
Circadian proteome of *C. reinhardtii*. **1A)** Schematic representation of the approach we used to capture the total proteome of *C. reinhardtii* across the two circadian cycles. The white and black rectangular boxes represent 12 hours of light and dark period respectively. The grey boxes represent the continuous light (LL) periods of subjective day and night respectively. Triplicate samples were collected from each time point for two respective cycles. **1B)** Venn diagram showing the number of common proteins found when we compared the LC-MS/MS yield from cycle 1 and cycle 2. **1C)** Principal component analysis (PCA) of the 1080 common proteins obtained from the two cycles. The two components PC1 and PC2, are represented ax X and Y-axis respectively. The day samples are shown here as red-filled circles and the night samples as filled blue circles. Each filled circle represents a replicate from the time-point (CT) mentioned in the figure. **1D)** Pie chart illustrating the percentage of cycling and non-cycling proteins out of the total 1080 proteins (from 1b) of *C. reinhardtii*.

The protein intensities across the biological and cycle replicates were normally distributed (Supplementary Figure 1C). Principal component analysis (PCA) (Figure 1C) and Pearson correlation plot (Supplementary Figure S1D) revealed a consistent time-dependent clustering of the biological and cycle replicates. Timewise clustering of samples showed an unambiguous distinction between the day and night samples, which are separated based on the significant component 1 represented on the X-axis (Figure 1C). Subjective dawn and midday samples cluster together, as do the subjective dusk and midnight samples. This suggests that the samples from different times of the day and night faithfully group with their temporal neighbours and the highest difference observed between dawn and dusk (Supplementary Figure 1E). Using the cosine fit function embedded in the MaxQuant Perseus analysis software (described in the M&M section) we identified that almost 48% (517/1080 proteins) out of the total captured proteome is oscillating with a free running period (Figure 1D). The remaining 563 proteins did not qualify as rhythmic across the two circadian cycles (Figure 1D). This is the first comprehensive study in *C. reinhardtii* that shows a significant number of proteins oscillate across the circadian cycle.

### The rhythmic proteome of *C. reinhardtii* unveils robust temporal segregation of protein accumulation

We further analysed the 517 rhythmic proteins to extract the phase distribution along the course of the circadian days. Using distribution and median line plotting, we found two distinct clusters, 291 proteins peak at CT54 and CT78 and trough at CT60 and CT84, corresponding to midday and dusk respectively. On the other hand, 226 proteins peak at dusk and are out-of-phase with the 291 proteins discussed above (Figure 2A). The change in protein intensity when cells transit from dark to light or vice versa is quite evident from the protein intensity distribution (Figure 2A) and the phase distribution rose plot (Figure 2B). In photosynthetic species, change from light-to-dark or vice versa is known as the transition zone, where the clock’s role to anticipate has been widely appreciated. Therefore, it is not uncommon for *C. reinhardtii* to demonstrate such dynamics at the protein level. Protein accumulates specifically at two times in *C. reinhardtii*, near midday and midnight respectively. Similar pattern of protein accumulation can be observed in an earlier diurnal proteomics study in *C. reinhardtii* (Supplementary Figure S2A-E) (Strenkert *et al*, 2019) and circadian proteomic studies in other species (Kay *et al*, 2021; Robles *et al*, 2014). Overall, our results clearly show the circadian rhythm of (291 + 226) 517 proteins, which contributes almost 48% of the total proteins we identified. Considering that we have identified only about 10% of the total proteins reported in *C. reinhardtii*, circadian regulation of protein accumulation seems to be widespread in this species.

**Figure 2:**
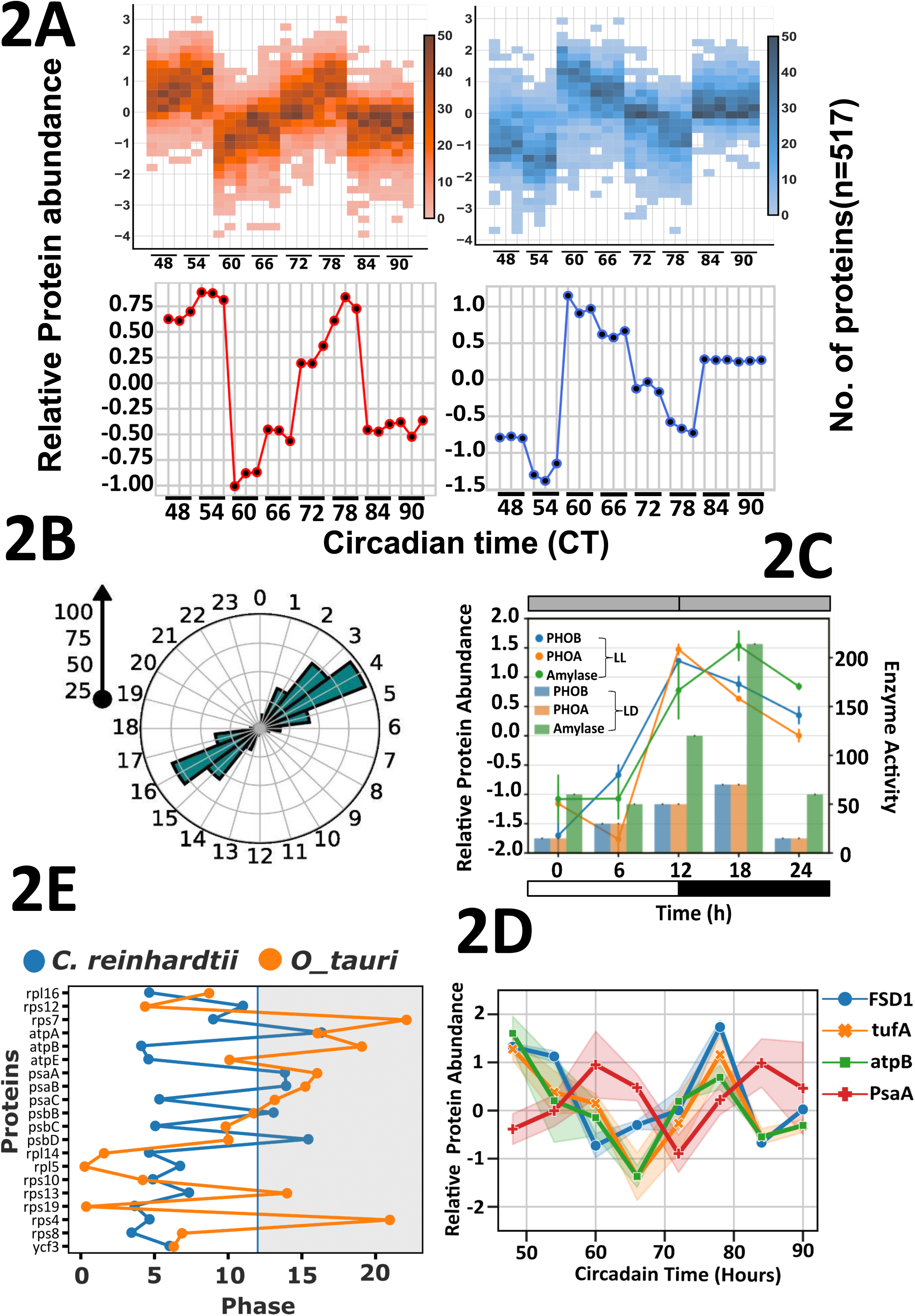
Daily dynamics of the circadian proteome. **2A)** Distribution plot revealing the significant day (top left) and night (top right) cluster of the cycling proteins (517). The calculated median of the proteins peaking during the day(red) and night(blue). **2B)** Rose plot presenting the phase distribution of the cycling proteins. The arrow on the top left represents the number of proteins. **2C)** Histogram depicting the daily changes in phosphorylase (**orange** PHOA, **blue** PHOB) and amylase enzyme activity (**green**). The overlapped line plot shows the relative protein abundance of phosphorylase PHOA (**orange** line), PHOB (**blue**), and amylase (**green**) the error bar stands for standard error. **2D)** Relative abundance of some proteins from *c.reinhardtii* showing robust day-night rhythms. **2E)** Line plot comparing the phase of organellar-encoded proteins (chloroplastic or mitochondrial) from previously studied *O. tauri* circadian proteome (**blue**) and our *C. reinhardtii* (**orange**) circadian proteome.

To verify that the SWATH LC-MS/MS approach and the downstream analysis portray the true nature of the proteome expression pattern in *C. reinhardtii*, we mined the published datasets to find evidence that would corroborate and validate our findings. First, we looked into Chlamy1, which is a non-oscillating RNA-binding clock protein (Zhao *et al*, 2004). Although Chlamy1 activity shows circadian changes, its immunoblotting study has confirmed its non-rhythmic nature across the circadian cycle. We used the same chlamy1 to confirm its circadian dynamics using western blotting. As expected, the Chlamy1 protein did not show any significant rhythm in the captured circadian proteome (Supplementary figure S3A). Our SWATH-MS analysis also corroborated this finding and showed no rhythm in chlamy1 dynamics across two cycles of circadian sampling (Supplementary Fig S3B). As a result, we believe that the SWATH proteome can provide an accurate measure of the non-rhythmic nature of this protein. Next, we compared two diurnally rhythmic proteins, ɑ-amylase and glycogen phosphorylase, whose diurnal activity profiles were revealed in an earlier study (Levi & Gibbs, 1984). Upon comparing the protein activity profiles we found the protein intensity dynamics of amylase and phosphorylase derived from our DIA-SWATH-MS analysis concur with the published data. Also, our analysis showed that these two proteins are regulated by the endogenous clock (Figure 2C). *tufA* is a well-known circadian clock protein (Hwang *et al*, 1996) and its protein intensity dynamics from our analysis agrees with its existing circadian profile from previous studies in *C reinhardtii* (Strenkert *et al*, 2019) and *Ostreococcus tauri* (Kay *et al*, 2021). Our circadian proteomics revealed *tufA* accumulation is circadian and its peak phase was around midday (Figure 2D). We also found 20 proteins from our analysis that are already known to oscillate with a circadian period in another green algae species, *Ostreococcus tauri* (Kay *et al*, 2021). When compared, we found that peak phases of 80% of these proteins are identical (Figure 2E). Our circadian phase distribution agrees with the diurnal protein distribution from an previous diurnal proteomics study and we found 47 proteins demonstrating identical protein accumulation patterns (Supplementary Figure S2A-H) (Strenkert *et al*, 2019). Accordingly, we conclude that the SWATH-MS and the subsequent downstream analysis successfully generated a considerable circadian proteome profile and its phase distribution pattern in *C. reinhardtii*.

### Circadian proteomics imparts molecular correlates of many known physiological rhythms in *C. reinhardtii*

We broadly categorized the 517 oscillating proteins into functional groups and mined their roles in generating rhythmic biological processes in *C. reinhardtii*. Classification of these proteins with gene ontology (GO) and KEGG analysis revealed many key pathways to be under circadian regulation. We grouped these processes as up at day or up at night (Figure 3A). The circadian proteomics directly portrays the daily dynamics of enzymes that are associated with key physiological pathways in *C. reinhardtii*. Some of the processes are unique and never have been reported before as circadian, while others are already verified as temporal based on the earlier findings, yet their circadian nature is revealed from our current analysis.

**Figure 3:**
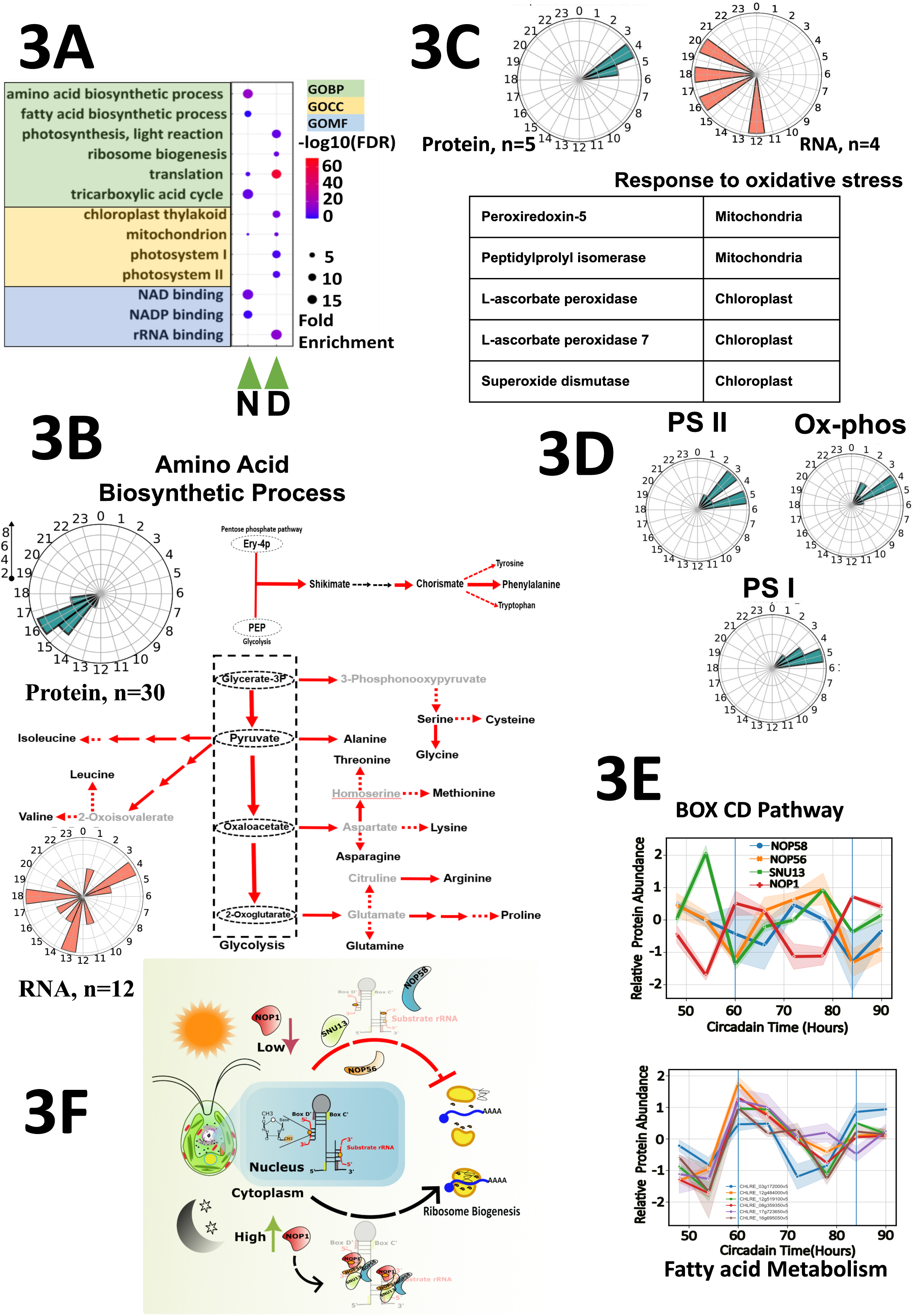
Circadian clock controlled cellular and metabolic pathways in *C. reinhardtii*. **3A)** Gene ontology-enriched categories (FDR <0.05) of the proteins associated day and night clusters. The diameter of the point represents fold enrichment and FDR is shown in color scale. Gene ontology biological process (GOBP) terms are highlighted in **green**, GO cellular component in **yellow**, and GO molecular function in **blue**. **3B) Left:** Rose plot illustrating the phase distribution of the cycling proteins (blue) and their encoding mRNAs (red) involved in the amino acid biosynthesis process. **Right:** Key enzymes involved in the amino acid biosynthesis process. **3C) Top:** Rose plot showing the phase distribution of proteins with antioxidant properties and their encoding mRNAs. **Bottom:** Table depicting the type of enzymes and their subcellular locations. **3D)** Phase distribution of proteins involved in photosystem I, photosystem II and oxidative phosphorylation **3D) Top panel:** Plot depicting dynamics of the proteins associated with BOX CD pathway across two subsequent circadian cycles. **Bottom panel:** Line plot showing the relative abundance of proteins involved in fatty acid metabolism. **3F)** A model depicting the role of 2’-O-methylation by BOX C/D complex in the regulation of global translation mediated by ribosomal translation efficiency.

*C. reinhardtii* is known to share conserved circadian elements with its distant kin *L. polyedra*. A recent circadian ribosome profiling study in *L. polyedra* showed a concerted expression of amino acid biosynthesis enzymes at around midnight (Bowazolo *et al*, 2022). Interestingly, our circadian proteomics study in *C. reinhardtii* also revealed a similar trend of amino acid biosynthesis at about the same phase, around midnight. About 30 enzymes belonging to the amino acid metabolic pathway peaked concurrently around CT15 to CT17 suggesting its coordinated regulation of protein accumulation by the clock (Figure 3B). The shikimate pathway connects carbohydrate metabolism to the synthesis of secondary metabolites and aromatic amino acids (Tzin & Galili, 2010). We also found the *C. reinhardtii* shikimate pathway to be controlled by the clock (Figure 3B). We next wanted to find out how many of these 30 rhythmic enzymes also demonstrate corresponding rhythm at the mRNA levels. We realized, firstly only about half of the 30 rhythmic enzymes have rhythmic mRNAs. Secondly, among the candidates where both mRNA proteins oscillate only a small fraction of them do have concurrent phase distribution. Lastly, half of the 30 rhythmic enzymes are generated from nonrhythmic mRNAs, thereby supporting the emerging notion of the widespread role of post-transcriptional machinery in clock regulation. Although the mRNAs involved in this pathway do not follow synchronized expression profiles, the protein accumulation is synchronized. Since amino acids are the building blocks necessary for protein synthesis to commence, an obvious question is does translation also follow a temporal regulation?

Subsequently, we found translation shows two peaks, the major peak is just between dawn and midday, where 47 of the total 197 ribosomal protein peaks concurrently (with fold enrichment of 30 and an FDR value of 4.5E-50) along with rate limiting translational initiation factor eIF2 and its regulator eIF2B. EIF5A also peaks during this time (Supplementary Fig S4A), which is known to play a crucial role in elongation and termination of the peptide chain (Schuller *et al*, 2017).

*C. reinhardtii* cells have a mechanism to resist UV damage during the day that phases out by the time cells transit to dusk (Nikaido & Johnson, 2000). Phase-dependent UV exposure of *C. reinhardtii* cells showed maximum resistance during the day that peaks around midday and gradually phases out by the end of the day (Nikaido & Johnson, 2000). Our proteomics analysis uncovered the biochemical players that possibly underlie this phenomenon. We found 5 proteins with antioxidant properties that are localised in the mitochondria and chloroplast and whose dynamics follow exactly the UV sensitivity pattern (Figure 3C) deduced from earlier studies. We also found that 4 out of these 5 proteins show rhythms in mRNA accumulation albeit their peak phases are randomly distributed across the night phase. It confirmed our earlier observation that the rhythm of mRNAs associated with a biological process may not be synchronized whereas their encoded protein rhythms are in perfect accord. This mismatch of rhythms between mRNA and its encoded proteins seems to be very widespread (Supplementary Figure S4B).

Furthermore, as expected, proteins primarily localized on the thylakoid membrane which are involved in regulating photosynthesis light reaction, and part of photosystem I and II peaks at midday (Figure 3D). Overall, our circadian proteome dynamics data authenticate many of the previously known biological rhythms and elaborate on the molecular players involved in these biological processes. Next, we used our circadian proteomics to identify additional circadian processes that remain elusive yet.

### Circadian proteomics reveals novel clock-driven processes in *C. reinhardtii*

Our chronological proteomics analysis revealed some novel biological processes to be governed by the clock that seems to be regulated post transcriptionally. Box C/D RNAs are non-coding RNAs and belong to the family of RNA-guided RNA modification systems widespread in eukaryotes and archaea but absent in prokaryotes (Matera *et al*, 2007). These RNAs form ribonucleoprotein structures (Box C/D RNPs) with putative proteins that catalyse the 2′-*O*-methylation of ribosomal RNAs, small nuclear (snRNAs), mRNAs and tRNAs at specific sites that are guided by the C/D RNAs (Kufel & Grzechnik, 2019). This modification can protect RNAs from getting cleaved, enhance codon recognition and their thermal stability, can act as chaperones, impact the global translational efficiency of the ribosomes, and modulate high-temperature folding (Therizols *et al*, 2015). In *C. reinhardtii* four proteins (NOP1, NOP56, NOP58, and SNU13) form the RNP complex with the box C/D RNAs to catalyse the methylation. We found all four proteins are rhythmic under circadian conditions (Figure 3E top panel). We found the NOP1 protein rhythm to be antiphase with the other 3 proteins involved in the BOX C/D pathway (Figure 3E top panel). NOP1 is the rate-limiting S-adenosyl-L-methionine-dependent methyltransferase that methylates RNAs as well as histone protein H2A (Tessarz *et al*, 2014) NOP1 mediated rRNA methylation can regulate the translational efficiency of ribosomes (Khoshnevis *et al*, 2022), which can play a key role in temporal regulation of global translational rates (Figure 3F).

Lipids are generated from fatty acids (FAs), which are molecules consisting of a diverse length of a hydrocarbon chain that contains different degrees of unsaturation. Lipids are essential biomolecules that cater to multiple cellular needs such as those required for membrane synthesis, serve as high-energy storage molecules, can be broken down to meet cellular energy demand, and can generate secondary signalling molecules (Okazaki & Saito, 2014). The first rate-limiting step of fatty acid biosynthesis is the conversion of Acetyl CoA to Malonyl CoA by a multi-subunit protein acetyl CoA-carboxylase (Davis *et al*, 2000). In *C. reinhardtii*, this enzyme has four subunits (Li-Beisson *et al*, 2015); we found 3 of the 4 subunits to have a circadian mode of expression. All these 3 subunits showed cyclical expression patterns peaking at night for the two subsequent circadian days that we tested (Figure 3E bottom panel). Another fatty acid biosynthesis enzyme 3-oxoacyl- [acyl-carrier protein] reductase, which most probably catalyses the first reduction step in the elongation cycle of FA biosynthesis (Li-Beisson *et al*, 2015), also peaks with the same phase at night. A previous study performed the metabolic analysis with day/night synchronized *C. reinhardtii* cells and showed a similar night-peaking pattern of many FAs (Zones *et al*, 2015; Strenkert *et al*, 2019).

Peroxisomal **β**-oxidation in *C. reinhardtii* is required for breaking down very long chain fatty acids (Kong *et al*, 2017). Jasmonic acid biosynthesis is a result of peroxisomal **β**-oxidation(Li *et al*, 2021). In *C. reinhardtii* jasmonic acid pathway is not complete and it does not go beyond 7 - iso-jasmonic acid CoA. The enzymes that catalyze the 7 - iso-jasmonic acid CoA to Jasmonic acid is not present in *C. reinhardtii*. The role of 7 - iso-jasmonic acid CoA is not known, however other family members of jasmonate play crucial roles in plant stress response and development (Li *et al*, 2021). We found two peroxisomal enzymes, namely acetyl-CoA acyltransferase 1 and enoyl-CoA hydratase/3-hydroxyacyl-CoA dehydrogenase that are regulated by the clock and are associated with peroxisomal **β**-oxidation of FAs that lead to the generation of 7 - iso-jasmonic acid CoA. The nightly peaking of these two enzymes suggests the synthesis of 7 - iso-jasmonic acid CoA or its derivatives at night. However, at this point we cannot comment about the physiological consequence as the role of 7 - iso-jasmonic acid CoA in *C. reinhardtii* is yet to be deciphered. These findings in *C. reinhardtii* are novel and might have important outcomes in future.

### Asynchrony in phase distribution of RNA and their encoded protein is prevalent in *C reinhardtii*

Recent studies have pointed out that the circadian clock can regulate protein turnover independent of the mRNA regulation (Robles *et al*, 2014) resulting in mRNA rhythms not corresponding to the rhythms of their encoded proteins across the daily cycle. From our observation in the preceding section, we realized that mismatch in mRNA and protein phase distribution can be rampant in *C. reinhardtii*. Hence, we decided to test the extent of this mismatch in *C. reinhardtii* using an already published diurnal transcriptome that uses time points corresponding to our study (Panchy *et al*, 2014). To be clear, the diurnal transcriptome provides both the diurnal and circadian information of transcripts across the 24-hour cycle. On the other hand, our data shows circadian protein abundance across the 24-hour cycle. By comparing these two datasets we can decouple candidates that are regulated by the environmental signal (light) from the ones controlled solely by an endogenous clock. We mined the transcriptome dataset to extract the mRNA phase information of 1080 genes whose protein phase information we got from this study. We found mRNA FPKM values of 1042 (out of 1080) candidates over the diurnal cycle. Using the cosine fit function, we evaluated the rhythmic nature of these 1042 transcripts. Based on their rhythmic nature, these 1042 candidates were divided into two groups, 1st group of 550 individuals whose proteins are non-rhythmic and 2nd group of 492 individuals whose proteins are rhythmic. Group 1a consists of 31% (170) of the 550 candidates whose mRNA and its encoded proteome are non-rhythmic (Figure 4A). Group 1b consists of the other 69% (380) of the 550 candidates, where the mRNA shows clear oscillation under a diurnal regime, although their encoded proteins showed no oscillation under a circadian cycle (Supplementary Fig S4C and S4D). We envisage two possibilities of this dysregulation between mRNA and protein oscillations. As the transcriptome is diurnal and the proteome is circadian, one possibility is light-dark regulation of transcription that does not influence the downstream protein synthesis. The other possibility is that these proteins do not reciprocate their parent mRNA oscillation, a characteristic that is prevalent in eukaryotes (Mauvoisin *et al*, 2014).

**Figure 4:**
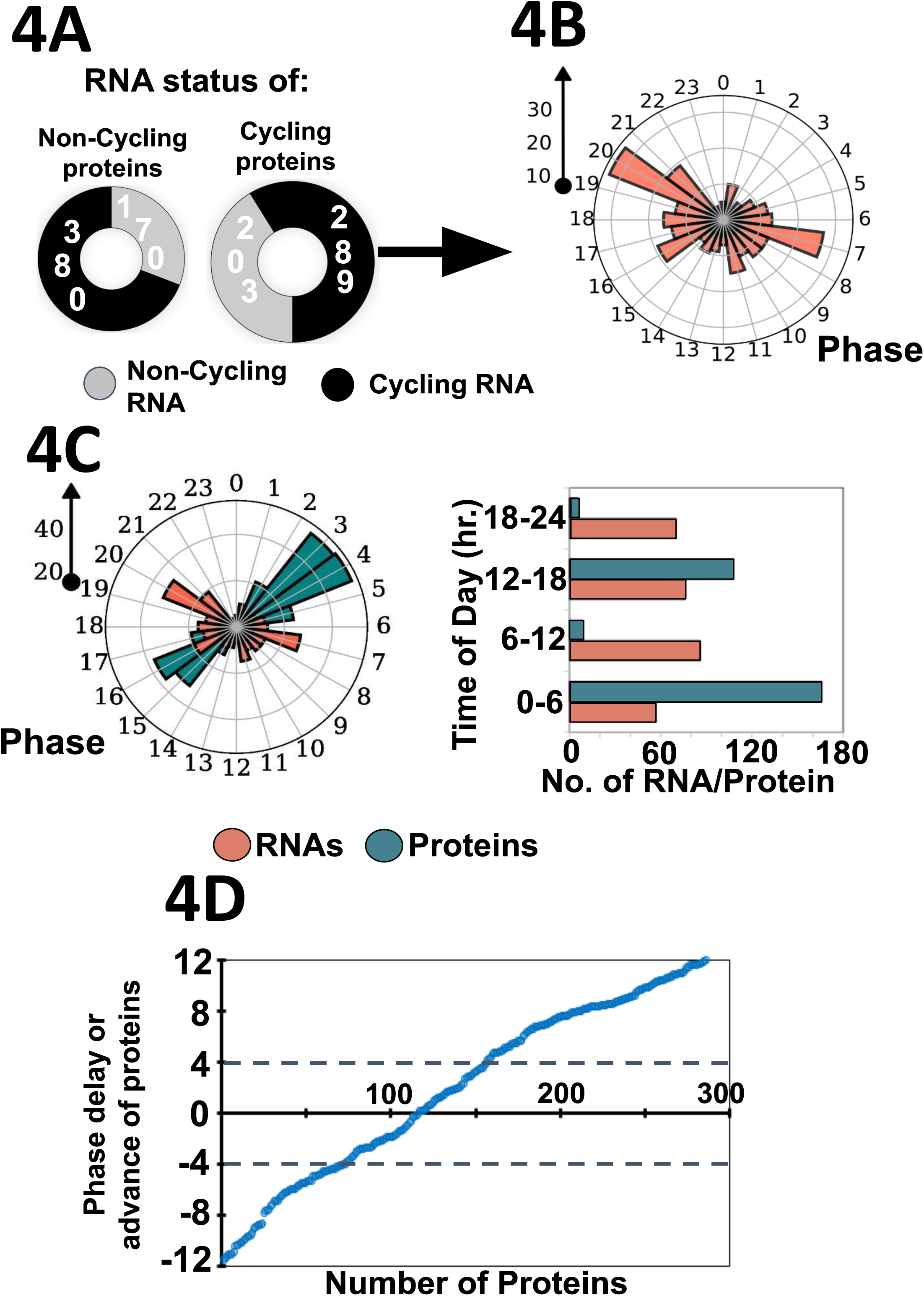
Phase distribution of mRNAs and their encoded proteins are not synchronous. **4A)** Pie chart of cycling proteins (left) with their cycling (colored **black)** and non-cycling (colored **grey**) RNAs and non-cycling proteins (right) their RNA with the same color scheme information. **4B)** Rose plot showing the phase distribution of the cycling mRNA whose encoded proteins (289) also cycles. Left arrow showing the number of RNAs **4C)** Left: rose plot with the phase distribution of the 289 cycling proteins (colored blue) overlapped with the phase distribution of the encoding mRNAs (colored red). Right: histogram showing the number of mRNAs (red) and protein s(blue) whose peak phase falls within the time shown in Y-axis. The time is binned in 6 hours. **4D)** Scatter plot revealing the phase difference between the mRNAs and proteins. Each dot represents a phase delay or advance of an individual protein from its respective parent mRNA (on the scale of 12 hours delay (−12) or advance (+12) on the Y-axis).

Group 2 (492 of 1042) candidates demonstrated robust protein rhythm across the circadian cycle. However, 41% (203 of 492) of these individuals showed no rhythms at their mRNA levels. We grouped them as 2a, these 203 are prospective candidates where clock regulation may occur at the post-transcriptional level (Figure 4A). Daily rhythms in proteins without any corresponding mRNA rhythms is a prevailing phenomenon in eukaryotes (Morse *et al*, 1989; Rey *et al*, 2018; Mauvoisin *et al*, 2014; Robles *et al*, 2014). In the other group 2b, 59% (289) of candidates are those whose mRNA and their encoded proteins oscillate across the circadian cycle (Supplementary Figure S4A and S4B). These group 2b (289) candidates provide an ideal platform to study the mRNA - protein phase synchronization across the circadian cycle. The mRNA peak phase distribution plot of 289 candidates shows even daily distribution across all phases with two modest peaks at midnight (LD20) and midday (LD7) (Figure 4B). Whereas we find widespread desynchronization when we compare the overall mRNA - protein oscillation profiles across the day (Figure 4C). We observed that while the majority (>90%) of the protein abundance spiked either early morning (CT0 - CT6) (~55% proteins) or early night (CT13 - CT18) (~35% proteins), their mRNAs peak-phase remains evenly distributed across the 24-hour cycle when binned in 6-hours (Figure 4C). To gain a deeper insight, we compared the peak phase of 289 (group 2b) mRNAs with their encoded proteins. Taking the mRNA peak as the reference we mapped the peak phase of its encoded protein on a +12-hour delay or −12-hour advance scale (Figure 4D). As our protein intensities are based on a 6-hour wide range we envisage that a delay or advance of 4 hours or less can be considered synchronous (where the mRNA and its encoded protein peak concurrently). Any delay or advance of the protein peak phase beyond that can be considered asynchronous (where mRNA and protein peak phase are not concurrent). We found that only 30 % of the individuals are synchronous, where mRNAs and their encoded proteins peak simultaneously. More than 65% (196 of 289) of the candidates are asynchronous, with significant phase differences. The peak phase of mRNA can even be completely out of phase to the peak phase of its encoded proteins. Altogether, comparing the transcriptome and the proteome data in the same system, we found that the phase distribution of the rhythmic mRNA transcriptome diverged prominently from the phases of the cycling proteome.

We found widespread disagreement between the phase distribution of mRNAs and their encoded proteins (Figure 3B, C, and 5). Intensity peaks of proteins associated with metabolic pathways are extremely synchronized, whereas the peak phase of their parent mRNAs is widely distributed across the 24-hour cycle. This observation holds true for many biological processes (Figure 5). In many cases we observed the protein peak is delayed by 4-6 hours from its parent mRNA peak, an observation that agrees with a previous study done with mammalian cell lines (Robles *et al*, 2014; Mauvoisin *et al*, 2014).

**Figure 5:**
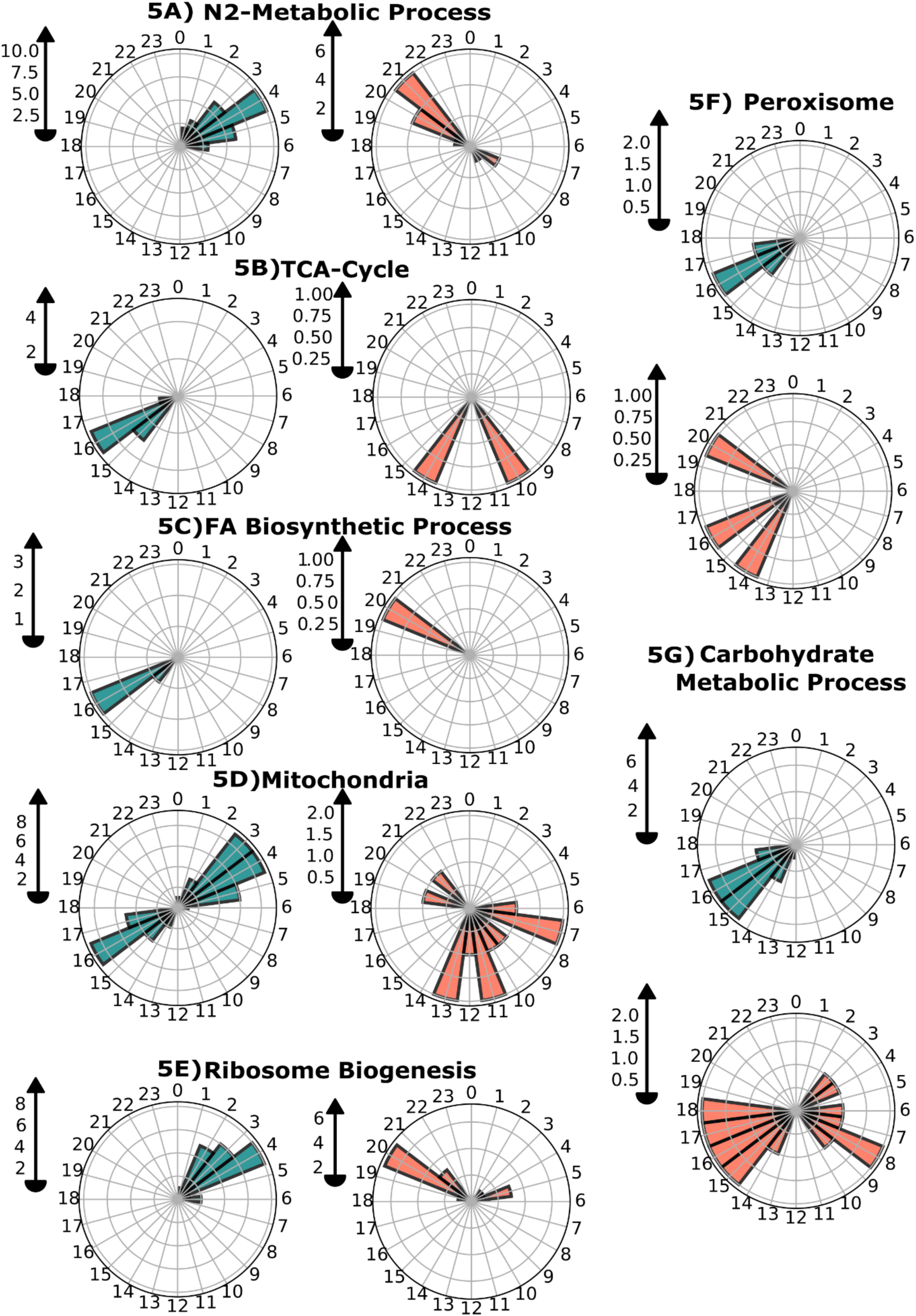
Circadian clock controlled cellular and metabolic pathways in *C.reinhardtii*. **5A-5L)** The left panel is an illustration of the phase distribution of proteins (blue-coloured rose plots) and the right panel shows the phase distribution of the respective encoding mRNAs (Tomato-coloured rose plots) associated with respective gene ontology (GO) categories that are mentioned on the top of each figure.

Proteins localized in mitochondria are required for aerobic respiration. Our analysis show some of these proteins accumulate during day and the others accumulate at night, this concurs with the mRNA rhythms observed in TCA cycle mediating transcripts (Zones *et al*, 2015). In photosynthetic plants, mitochondrial metabolic processes are known to differ between day and night (Lee *et al*, 2010). This is largely driven by the daily regulation of metabolism that feeds mitochondria with different substrates depending on the time of the day or night.

## Discussion

Genomic and transcriptomic studies were used as gold standard technologies to understand the global circadian regulation. Role of protein abundance in temporal regulation of physiology and metabolism has been underestimated (Takahashi, 2017a). Lack of in-depth and comprehensive quantitative proteomics in circadian biology were in part due to technological challenges in mass spectrometry approaches (Michalski *et al*, 2011; Tabb *et al*, 2010; Bondarenko *et al*, 2002; Zhu *et al*, 2010). High-resolution mass spectrometry like SWATH-MS not only identifies but also accurately quantifies thousands of label-free proteins across samples with high reproducibility (Collins *et al*, 2013, 2017). Thus, providing an ideal platform to reveal the proteome-wide dynamics across the daily cycle. These new features of mass spectrometry revolutionized the field of circadian proteomics providing a deeper insight, verifying, and complementing the mRNA level changes to their corresponding protein levels.

In *C. reinhardtii*, predominantly transcriptomic studies formed the basis to realize the clock driven physiological rhythms. Nearly 80-85% of the *C. reinhardtii* transcriptome shows diurnal rhythmicity (Strenkert *et al*, 2019; Zones *et al*, 2015). Clocks can attune post transcriptional pathways to accomplish daily rhythms without affecting the mRNA abundance (Morse *et al*, 1989; Roy *et al*, 2014; Mauvoisin & Gachon, 2020; Mauvoisin *et al*, 2014; Robles *et al*, 2014; Chiang *et al*, 2014; Kay *et al*, 2021; Takahashi, 2017a). Although evidence suggests an extensive role of post transcriptional machinery in clock driven cellular physiology of *C. reinhardtii* (Wagner *et al*, 2004, 2005; Wagner & Mittag, 2009), comprehensive studies in this direction are lacking. Our study revealed that almost 48% of the measured proteins to be rhythmic under circadian conditions (Figure 1). This is significantly higher than the proteome-wide rhythmicity observed in higher organisms like Arabidopsis, where it ranges from 0.4% to 10% depending on the methodology employed (Choudhary *et al*, 2016; Krahmer *et al*, 2021). Mammalian hepatic and liver proteomes showed 20% and 6% of the measured proteome to be rhythmic (Reddy *et al*, 2006; Mauvoisin *et al*, 2014; Robles *et al*, 2014; Chiang *et al*, 2014; Wang *et al*, 2018). On the other hand, 7% and 25% of the quantified proteins were rhythmic in Drosophila and *N. crassa* respectively (Rey *et al*, 2018; Hurley *et al*, 2018). Another interesting feature is the peak phase distribution of the rhythmic proteins in *C. reinhadrtii*. We found almost >95% of the rhythmic proteins either peak around midday or around midnight (Figure 2A, B). Similar observations concur with reports from other mammalian cells (Hurley *et al*, 2018; Rey *et al*, 2018; Mauvoisin *et al*, 2014; Robles *et al*, 2014). Similar phase distributions of protein accumulation was observed in an earlier diurnal proteomics study in *C. reinhardtii* (Supplementary Figure 2B-E) (Strenkert *et al*, 2019). Besides in Arabidopsis (Missra *et al*, 2015) and U2OS cell lines (Jang *et al*, 2015) synchronized with dexamethasone, where translation peaks around midday and midnight. The rhythmic proteins have a lower amplitude than the cycling mRNAs (Supplementary Figure 6), a feature that agrees with all circadian or diurnal proteomics studies (Bowazolo *et al*, 2019; Mauvoisin *et al*, 2014; Robles *et al*, 2014). Two plausible reasons can justify our observation, one is that protein turnover rates are slower than mRNAs. Secondly, to capture the true circadian rhythms we allowed cells to free run for 48 hours in constant condition (LL) before collection of cycle 1 samples. The damping of rhythms in free running conditions can also contribute to the damped oscillations in our case. We observed cycle 2 amplitudes are clearly more damped than cycle 1 because it underwent another round of free running.

This study reveals the pervasiveness of clock regulation of *C. reinhardtii*’s post-transcriptional machinery that generates rhythmic accumulation of proteins associated to key physiological and metabolic pathways (Figure 3, Supplementary figure 4B and S7). 41% of the rhythmic proteome in *C. reinhardtii* are generated from non-rhythmic mRNAs (Figure 4). These are prospective targets of post-transcriptional regulation where protein synthesis and /or degradation rather than mRNA abundance is under clock regulation. Different studies with mice showed almost 5% to 10% of the proteins are rhythmic and 20% to 50% of the rhythmic proteome do not show corresponding rhythms in mRNAs (Mauvoisin *et al*, 2014; Robles *et al*, 2014; Chiang *et al*, 2014; Azimifar *et al*, 2014; Wang *et al*, 2018). This mismatch is even higher in Drosophila (Rey *et al*, 2018) and the fungus *N. crassa* (Hurley *et al*, 2018). The notion that the mRNA levels neither correlate with the levels of their encoded proteins nor with their activity (Roy *et al*, 2014; Morse *et al*, 1989; Robles *et al*, 2014) is widespread. 59% of the rhythmic proteins in *C. reinhardtii* also showed rhythms in their parent mRNAs (Figure 4). But do they share the same phase distribution pattern? Our analysis showed a major mismatch between the rhythms of the mRNAs and their encoded proteins (Figure 4, 5 and Supplementary Figure 4B). This mismatch can be to such an extent that the two rhythms render a completely out-of-phase relationship (Figure 4D). Therefore, it is rational that protein rhythms should be used to complement the transcriptome-wide studies while predicting the phase of physiological rhythms. The remarkable coordination of protein levels associated with key cellular pathways suggests its key role in temporal regulation of diverse biological pathways in *C. reinhardtii* (Figure 5, Supplementary Figure S4B and S7). In *C. reinhardtii* substantial portion of rhythmic proteome arises from nonrhythmic mRNAs. In majority of the cases, we find the rhythmic proteome to be highly synchronized whereas its encoding rhythmic mRNAs peak phases are widely distributed across the day/night cycle. Apart from the transcription, protein synthesis and degradation as well as protein secretion contributes significantly generating rhythms at the proteome level. It is plausible that all these steps are independently regulated by the inherent clock that ultimately leads to a mismatch between mRNA and protein rhythms.

*C. reinhardtii* cells must protect themselves from UV using different strategies. One way can be circadian programming of UV sensitive tasks at night with the help of an endogenous oscillator (Nikaido & Johnson, 2000). Another way can be to regulate the cellular antioxidant system to counter for UV-mediated oxidative stress. Our circadian proteomics provides the molecular correlates of this UV protective mechanism during the day. Proteins functioning as antioxidants follow a circadian pattern of expression that correlates with the earlier observation of UV sensitivity pattern. The antioxidant proteins peak collectively during the day whereas their mRNAs phase distribution shows significant mismatch (Figure 3C). Protein intensity changes in our analysis alone may not be able to explain the activity of UV protection entirely, however, clocks can also modulate these enzymes allosterically and if they act in tandem with their protein intensity, it can create a greater impact in protecting the cells during the day. Similar oscillations of antioxidant enzyme activity are not uncommon in unicellular alga.

2’-O-methylation is a regular and abundant modification of coding and non-coding RNAs (Lapinaite *et al*, 2013). This methylation is known to be catalyzed by fibrillarin (NOP1 in *C. reinhardtii*) and its specificity is guided by a snoRNA-protein complex, where proteins NOP56, NOP58 and SNU13 play crucial roles. Our study found that all 4 proteins involved in this process are rhythmic under LL (Figure 3E). Fibrillarin downregulation impairs ribosomes resulting in loss of translational efficiency whereas its upregulation enhances ribosomes translational capabilities (Erales *et al*, 2017; Khoshnevis *et al*, 2022). Our study inculcates the possibility of temporal regulation of ribosomes ability to translate through rhythmic expression of the rRNA 2’-O-methylation machinery (Figure 3F) in *C. reinhardtii*. Although, at this stage it is just a possibility which remains to be validated with further studies. However, this mode of global regulation of translation by the clock has not been reported before (Erales *et al*, 2017; Khoshnevis *et al*, 2022) and can be an interesting topic of future investigation.

External light and the endogenous circadian clock coordinate with each other to modulate the daily dynamics of metabolism and biochemistry in photosynthetic species. A plethora of diurnal transcriptome studies were employed to understand this temporal regulation of physiology in *C. reinhardtii*. However, decoupling the clock’s contribution in these daily physiological dynamics is limited. Considering that proteins are the mediators of biological functions, proteome-wide studies to understand the circadian dynamics of proteins are lacking in this species. Our circadian proteome analysis revealed rampant regulation of post-transcriptional machinery by the clock to generate daily rhythms in key metabolic output. We envisage that in conjunction with the existing transcriptome dynamics data our protein intensity statistics will refine the mechanistic understanding of the circadian regulation of biological processes in *C. reinhardtii*. *C. reinhardtii* has links to the last common ancestor of plant and animal, therefore, understanding the temporal control of physiology and metabolism in this species will be the key to reveal the evolution of clock regulation across kingdoms.

## Materials and Methods

### Cell Culture and sample collection

*Chlamydomonas reinhardtii (CC125)* cells (A kind gift from Prof. B. J. Rao, IISER Tirupati, originally obtained from the *Chlamycollection.org*) were maintained at 12:12 hours of light: dark (LD) regime at 18 ± 1 °C on TAP agar plates supplemented with 100 µg/ml carbenicillin and 40 µg/ml carbendazim in a Percival algal growth chamber (AL41L4) with 75 µmoles m^−2^ s^−1^ light intensity. From three plates, loop full of *C. reinhardtii* cells were inoculated in three 100 ml Erlenmeyer flasks containing 40 ml of Tris-acetate-phosphate (TAP) liquid media plus 100 µg/ml carbenicillin and entrained under 12:12 hours LD cycle. Once the OD_680_ reached 0.4, cells were transferred into three separate 1000 ml Erlenmeyer flasks containing 400 ml of liquid TAP media. The cells were maintained in a 12:12 LD cycle for 5 days to synchronize the circadian clock. Subsequently, the light regime was changed to a 12:12 light-light (LL) to allow the cells to run on their free-running period. Cells were grown in LL conditions for two days. On the third day, cells were collected from the 3 different flasks (independent biological replicates) at CT_48, CT_54, CT_60, CT_66, CT_72, CT_78, CT_84, and CT_90 by centrifugation at 5000 rcf at 4 °C for 10 min. To keep the cell number constant across the time, OD_680_ (Strenkert *et al*, 2019) was measured before each collection and the volume was adjusted to approximately 40 million cells. The cells acquired from CT-48 to CT-66 were labelled as cycle 1 whereas those collected from CT-72 to CT-90 were labelled as cycle 2. Pelleted cells were either used fresh or stored at −80 °C until further use.

### Protein Purification using Acidic-Guanidinium-Thiocyanate-Phenol-Chloroform (AGPC)

For total protein purification, cell pellets were resuspended in lysis buffer [10mM Tris-HCL pH 7.4, 500mM NaCl, 1mM EDTA pH8, 5mM DTT, 80U RNase out (Invitrogen cat no 10777-019) and complete protease inhibitor cocktail (Sigma cat no P9599)]. After resuspension, cells were lysed at 19 kPa using a cell disruption one-shot (OS) system (Constant Systems LTD). 100 µL of the cell lysate (equivalent to approximately 40 million cells) were transferred to 1 ml PureZOL reagent (BIO-RAD cat no 7326890) and thoroughly mixed by shaking for 15 seconds, later incubated at room temperature for 5 minutes. For the phase separation, 200 µl of chloroform was added to the reaction, mixed vigorously for 15 seconds, and incubated for 5 min at room temperature. After the incubation, samples were centrifuged at 15,000 rcf for 10 min at 4 °C. The lower organic phase was transferred into a fresh tube, proteins were precipitated with 100 (%) ethanol. The precipitated protein samples were resuspended in 100 µl of 100mM triethylammonium bicarbonate (TEAB) buffer pH 8.5 (Sigma-Aldrich cat no - T7408).

### Sample Preparation for SWATH-MS Proteomics

#### i) Reduction, Alkylation, and Trypsin Digestion

The Equivalent concentration of protein from each sample was reduced with 10 mM of Dithiothreitol (DTT) for 45 minutes at 56 °C, followed by alkylation using 20 mM of Iodoacetamide (IAA) at room temperature (in the dark) for 45 minutes as per the protocol described elsewhere(Villanueva *et al*, 2020) These samples were then subjected to trypsin (*cat no* V5111 sequencing grade, Promega) digestion in an enzyme to substrate ratio of 1:20 (trypsin: protein) for 16 hours at 37 °C at 400 rpm. Finally, the reaction was quenched by adding 0.5% TFA. Peptide desalting was performed using C18 spin columns (Pierce *cat no-89851)* as per the manufacturer’s protocol. Peptide quantification of each sample was performed using Bradford. The tryptic peptides were vacuum dried in a vacuum concentrator (Eppendorf) and each sample was cleaned up using Oasis HLB 1 cc Vac cartridges (Waters) using the manufacturer’s protocol. For the preparation of the spectral ion library, 300 µg of protein from all the samples were pooled and digested using trypsin as described above.

#### ii) Spectral ion library generation

Tryptic peptides from the protein pool were fractionated into 8 fractions by cation exchange cartridge (part no. 4326695) and using an increasing concentration of ammonium formate buffer (35 mM-350 mM ammonium formate, 30% v/v ACN, and 0.1% formic acid; pH = 2.9). Peptides from each of these fractions were cleaned up using C18 ZipTip (Millipore, USA). Each fraction was then analysed on a quadrupole-TOF hybrid mass spectrometer (TripleTOF 6600, SCIEX) coupled to an Eksigent NanoLC-425 system. Optimized source parameters were used, curtain gas and nebulizer gas were maintained at 25 psi and 20 psi respectively, the ion spray voltage was set to 5.5 kV and the temperature was set to 250 °C. About 4 μg of peptides were loaded on a trap column (ChromXP C18CL 5 µm 120 Å, Eksigent, SCIEX) and online desalting was performed with a flow rate of 10 µl per minute for 10 min. Peptides were separated on a reverse-phase C18 analytical column (ChromXP C18, 3 µm 120 Å, Eksigent, SCIEX) in a 57-minute-long buffer gradient with a flow rate of 5 µl/minute using water with 0.1% formic acid (buffer A) and acetonitrile with 0.1% formic acid (buffer B) as follows:

| **Time (min)** | **% A** | **% B** |
| --- | --- | --- |
| **0** | 97 | 3 |
| **38** | 75 | 25 |
| **43** | 68 | 32 |
| **45** | 20 | 80 |
| **45.5** | 10 | 90 |
| **48** | 10 | 90 |
| **49** | 97 | 3 |
| **57** | 97 | 3 |

Data was acquired using Analyst TF 1.7.1 Software (SCIEX). A 1.3-sec instrument cycle was repeated in high sensitivity mode throughout the entire gradient, consisting of a full scan MS spectrum (400–1250 m/z) with an accumulated time of 0.25s, followed by 20 MS/MS experiments (100–1500 m/z) with 50 msec accumulation time each, on MS precursors with charge state 2+ to 5+ exceeding a 120-cps threshold. The rolling collision energy was used, and the former target ions were excluded for 15 seconds.

#### iii) SWATH-MS data acquisition

The in-solution digested samples from the two cycles were analysed in SWATH-MS mode on the same instrument with similar LC gradient and source parameters as DDA runs. A SWATH-MS method was created with 100 precursor isolation windows, defined based on precursor m/z frequencies in DDA run using the SWATH Variable Window Calculator (SCIEX), with a minimum window of 5 m/z. Accumulation time was set to 250 msec for the MS scan (400– 1250 m/z) and 25 msec for the MS/MS scans (100–1500 m/z). Rolling collision energies were applied for each window based on the m/z range of each SWATH and a charge 2+ ion, with a collision energy spread of 5. The total cycle time was 2.8 sec.

### Mass Spectrometric Data analysis

A merged database search for DDA runs was performed using Proteinpilot™ Software 5.0.1 (SCIEX) against *Chlamydomonas reinhardtii* proteome from UniProtKB (UP000006906, with 18,829 protein entries). The Paragon algorithm was used to get protein group identities. The search parameters were set as follows: sample type-identification, cysteine alkylation-iodoacetamide, and digestion-trypsin. The biological modification was enabled in ID focus. The search effort was set to ‘Thorough ID’ and the detected protein threshold [Unused ProtScore (Conf)] was set to >0.05 (10.0%). False discovery rate (FDR) analysis was enabled. Only proteins identified with 1% global FDR were considered true identification. The group search result file from Proteinpilot™ Software was used as a spectral ion library for SWATH analysis. SWATH peak areas were extracted using SWATH 2.0 microapp in PeakView 2.2 software (SCIEX), and shared peptides were excluded. SWATH run files were added and retention time calibration was performed using peptides from abundant proteins. The processing settings for peak extraction were a maximum of 10 peptides per protein, 5 transitions per peptide, >95% peptide confidence threshold, and 1% peptide FDR. XIC extraction window was set to 5 min with 50 ppm XIC Width. All information was exported in the form of MarkerView (.mrkvw) files. In MarkerView 1.2.1 (SCIEX), protein area data was normalized, the data normalization strategy used was total area sum normalization and further analysis was performed in a Microsoft Excel file.

### Bioinformatics and Statistical Analysis

#### i. Batch Correction

Cells from the two cycles were collected and processed simultaneously. However, the LC-MS/MS sequencing of the two cycles were done in two batches. Cycle 1 and cycle 2 replicates were sequenced separately in two different batches 1 month apart, in the same mass spectrometry facility and with the same mass spectrometer machine. Subsequently, to remove any difference caused due to non-biological reasons, we performed “ComBat”, a batch correction software function embedded in BatchServer (Zhu *et al*, 2021) with default parameters on common proteins identified from cycle 1 and cycle 2. The PCA plots were generated using the Perseus software (Tyanova *et al*, 2016). All the further analysis was done with the batch-corrected datasets.

#### ii. Parameters for identification of Circadian Cycling Proteins

To identify the cycling proteins from cycle 1 and cycle 2, the protein abundance profile was fitted in a cosine (embedded in Perseus software) with a fixed period of 23.6 hours and the amplitude and phase were set as the free parameters. To visualize the cycling proteins, Z-normalised relative protein intensities from all the time points were visualized using the density. The density plot and the rose plot were created using the seaborn package in Python version 3.9.13.(https://www.python.org)

#### iii. Parameters for Diurnal Cycling Proteins

Publicly available proteome data at ZT0, ZT5, ZT11, ZT13, ZT18 and ZT23 time points of *Chlamydomonas reinhardtii,* original study (Strenkert *et al*, 2019) was obtained. Proteins with minimum 1 peptide at selected time points were used for further analysis. The Z normalized protein abundance was fitted in cosine with the fixed period of 24 hours. In addition, amplitude and phase were set as free parameters. Further diurnal cycling proteins were compared with the circadian cycling proteins using the gene ID. Density plots and phase distribution plots were created using the python mentioned in circadian cycling section.

#### iv. Parameters for identification of Diurnal cycling mRNA

Diurnal transcriptome data of *Chlamydomonas reinhardtii* was downloaded from the gene expression atlas ([Experiment < Expression Atlas < EMBL-EBI](https://www.ebi.ac.uk/gxa/experiments/E-GEOD-62671/Results)) original study(Panchy *et al*, 2014). The mean FPKM values of the common RNA were log2 transformed and the RNA abundance profile was fitted in a cosine with a fixed period of 23.6 hours using MaxQant Perseus software. The phase distribution of the cycling RNA was visualized using a rose plot created using python.

#### v. Gene Ontology KEGG pathway Analysis

Gene enrichment analysis of total identified day and night abundant proteins was performed using PANTHER V17.0(Mi *et al*, 2019) (http://www.pantherdb.org/). The enrichment was done using the default Fisher exact test with Benjamini-Hochberg FDR correction. A p-value cut-off of 0.05 was used to filter nonsignificant GO-terms. The fold enrichment and the FDR were visualized either using R or Python. KEGG pathway analysis was done using the KEGG mapper (Kanehisa & Sato, 2020)

### Western Blotting and SDS PAGE

Equal concentrations of protein crude extract from 4 different time points (CT48, CT54, CT60, and CT66) were separated on 12% SDS-PAGE. The proteins were transferred onto the PVDF membrane using a semi-dry transfer apparatus. Transfer quality was assessed by using ponseas-S stain. After the transfer, the membrane was blocked in 3% BSA for 1 hour at room temperature. Histone H3 antibody (Agrisera) and C3 (a kind gift from Professor Maria Mittag was diluted in 3 % BSA with 1:5000 dilution. The blot was incubated with primary antibodies overnight at 4 °C. Next day, the membrane was rinsed 3 times with TBST as per the standard western blotting protocol. Goat anti-rabbit antibody was diluted in 3% BSA (1:100,000) and the blot was incubated with a secondary antibody for 1 hour at room temperature. The blot was rinsed three times with TBST. Finally, the blot was developed using a chemiluminescence substrate as per the manufacturer’s protocol. The image was captured in the Chemi-Doc (BIO-RAD) instrument using default parameters. The band intensity was calculated using open-source ImageJ software.

## Data availability

The mass spectrometry proteomics raw data have been deposited to the ProteomeXchange Consortium via the PRIDE (Vizcaíno *et al*, 2016) partner repository with the dataset identifier PXD043839. Along with the raw data we have also uploaded the excel files with all details.

## Acknowledgements

The authors thank Dr. Shantanu Sengupta and Mr. Praveen Singh of the National Facility for Biochemical and Genomic Resources (**NFBGR**), IGIB for help with SWATH-MS sequencing. We thank Prof. Maria Mittag for providing us the Chlamy 1 antibody. This study was supported by the Science and Engineering Research Board, Government of India (SERB grant SRG/2019/000364), SR acknowledges the Annual Research Grant from Ashoka University. DBJ was supported by a PhD fellowship from Ashoka University.

## Author contributions

**Dinesh Balasaheb Jadhav:** Conceptualization; data curation; software; experimentation - designing and performing; formal analysis; validation; investigation; visualization; methodology; writing – initial draft, review and editing. **Sougata Roy:** Conceptualization; experiment designing; resources; supervision; project administration; formal analysis-some: funding acquisition; writing – original draft; writing – review and editing.

## Supporting Information

Supplementary Figures (PDF)

